# Defining the role of aerobic respiration in the metabolism and bioenergetics of *Enterococcus faecalis*

**DOI:** 10.64898/2026.08.30.748090

**Authors:** Olivia Paxie, Brunda Nijagal, Francesca O Todd Rose, Samuel Gastrell, Soleil Su, Anas Saleh, James W Grimshaw, Kyu Rhee, Henrik Strahl, Gregory M Cook, Rachel L Darnell

**Affiliations:** Department of Microbiology and Immunology, University of Otago, Dunedin, New Zealand; Metabolomics Australia, Bio21 Molecular Science and Biotechnology Institute, Parkville, Australia; Department of Medicine, Weill Cornell Medicine, New York, New York, USA; Centre for Bacterial Cell Biology, Biosciences Institute, Faculty of Medical Sciences, Newcastle University, Newcastle upon Tyne, UK

**Keywords:** Electron transport chain, NADH oxidase, energy metabolism, *Enterococcus*, bioenergetics, proton motive force, membrane potential

## Abstract

*Enterococcus faecalis* is an opportunistic pathogen and facultative anaerobe that primarily relies on fermentative metabolism to colonize a wide range of aerobic and anaerobic environments. In the presence of exogenous heme, *E. faecalis* can assemble a minimal electron transport chain consisting of membrane-associated primary dehydrogenases, demethylmenaquinone, and the terminal cytochrome *bd* oxidase (CydAB). This respiratory chain is thought to generate a proton motive force to drive ATP synthesis via the F-type ATP synthase, thereby improving energy conservation under aerobic conditions. However, a cytosolic NADH oxidase (Nox) also consumes NADH and oxygen, potentially competing with the electron transport chain for reducing equivalents and terminal electron acceptors; but the relative physiological contributions of these two oxygen-reducing pathways remain poorly understood. To define the roles of CydAB and Nox under normoxic and hypoxic conditions, we constructed Δ*cydAB* and Δ*nox* mutants. Real-time, *in situ* measurements revealed Δ*cydAB* had no significant effect on oxygen utilization while in the Δ*nox* it was significantly reduced; revealing Nox as the major consumer of oxygen. Semi-untargeted metabolomic analysis further revealed oxidase-specific alterations in central metabolism with the Δ*nox* causing pronounced shifts in the ATP and NADH ratios; highlighting Nox as a key determinant of intracellular redox and energy homeostasis. Finally, single-cell fluorescence microscopy showed that membrane potential, a component of proton motive force, was substantially diminished only in the absence of both CydAB and Nox, or the F-type ATP synthase. These findings indicate that the F-type ATP synthase is a major generator of proton motive force, even upon aerobic growth, and demonstrate a complementary role for the electron transport chain and Nox in the bioenergetics of *E. faecalis*.

## Introduction

Enterococci are facultative anaerobes capable of occupying anaerobic intestinal habitats while also surviving and dispersing through aerobic environmental reservoirs such as soil, water, plants, food systems, and hospital surfaces (1–5). *Enterococcus faecalis* features a diverse array of metabolic pathways to facilitate fermentation including glycolysis and the pentose phosphate pathway, which reflects its native environment, the nutrient-competitive, microoxic gastrointestinal tract (3,6). Under anaerobic growth, its primary fermentation product is L-lactate but depending on growth conditions, redox balance, and carbon source, formate, ethanol and acetate can be produced as secondary mixed-acid fermentation products (7,8). However, despite the breadth of fermentative metabolic pathways, *E. faecalis* is able to respire aerobically when grown in the presence of exogenous heme (9–11).

The predicted components of the aerobic electron transport chain (ETC) in *E. faecalis* include a membrane-bound type II NADH dehydrogenase (Ndh2; EF2055), which allows for re-oxidation of NADH, and a non-proton pumping cytochrome *bd* terminal oxidase (CydAB; EF2061 – 2060) (10). Heme enables the reconstitution of an active CydAB to reduce oxygen to water while potentially contributing to a proton motive force (PMF) through the generation of ΔpH via scalar protolytic reactions (12). As *E. faecalis* is unable to synthesise heme itself, they rely on heme uptake via the heme ABC transporter CydCD (EF2059 – 2058) encoded in the operon with CydAB (13). These two processes are coupled via demethylmenaquinone (DMK), which serves as a mobile electron carrier between electron-donating (NADH oxidation) and electron-accepting (oxygen reduction) reactions (14). ATP production is postulated to occur via the F-type ATP synthase (ATP synthase; EF2614 – 2607), driven by the PMF generated at least to a degree by the ETC. However, earlier studies in other enterococcal species argue for an alternative role of the ATP synthase as an ATP-driven proton pump involved in pH homeostasis (15–17).

Genes encoding for this basic ETC are also found in other enterococci including *E. cassiflavus, E. gallinarium,* and *E. avium* but, intriguingly, all are absent in the genomes of *E. faecium* and *E. hirae* (18). The presence of an active ETC has been linked to increased ATP levels and bacterial biomass yields in closely related lactococci and streptococci, thus arguing for a significant positive contribution towards the cell’s metabolic efficiency (19–23). However, no studies have systematically investigated their role in the bioenergetics of *E. faecalis*, and this metabolic pathway thus remains poorly understood.

In the absence of an efficient ETC many lactic acid bacteria possess cytoplasmic NADH oxidases, which are functionally distinct from the membrane-bound respiratory oxidases (24). However, Nox-like enzymes have been shown to promote aerobic growth in closely related streptococci including *S. pyogenes, S. mutans,* and *S. pneumoniae* (25–27). Under aerobic conditions, NADH oxidase allows for oxygen consumption and the reoxidation of NADH (28). *E. faecalis* encodes a water-forming type 2 NADH oxidase (Nox; EF1586), but its contribution to the bioenergetics of *E. faecalis* is currently unknown (27).

While oxygen consumption and energy generation by the ETC and accessory oxidases is well established and understood, comparatively little is known about how multiple terminal oxidases function together in facultative anaerobes. The uncommon dual-oxidase system of *E. faecalis* therefore provides a valuable model for understanding the functional coordination of respiratory enzymes and their contributions to metabolic flexibility. To investigate the metabolic and bioenergetic contributions of molecular oxygen in *E. faecalis*, we generated gene deletion mutants deficient in the cytoplasmic NADH oxidase and the aerobic ETC components cytochrome *bd* oxidase and F-type ATP synthase. Real-time measurements of oxygen consumption, semi-untargeted metabolomic profiling, and assessments of membrane potential were used to define the individual contributions of these enzymes to oxygen-dependent metabolism and cellular bioenergetics.

## Results

### Aerobic respiration contributes to the bioenergetic efficiency of *E. faecalis*

To determine the role of aerobic respiration in *E. faecalis* growth, we carried out growth assays of the wild-type (JH2-2) strain in heme-containing, nutrient-rich media. Two different oxygen tension (normoxia and hypoxia) were used to assess the bioenergetic contribution of aerobic respiration. Normoxia is defined as cultures grown upon shaking in conical flasks that allow free oxygen diffusion, providing an initial oxygen tension of 45 % (Fig. S1A). Hypoxia, in contrast, was achieved by culturing upon shaking in sterile, stoppered serum vials that do not allow free oxygen diffusion, providing an initial oxygen tension of 25 %, dropping to below 1 % after 5 h (Fig. S1B). The wild-type cells grew comparably in both oxygen-defined conditions (Fig. 1).

**Fig. 1.**
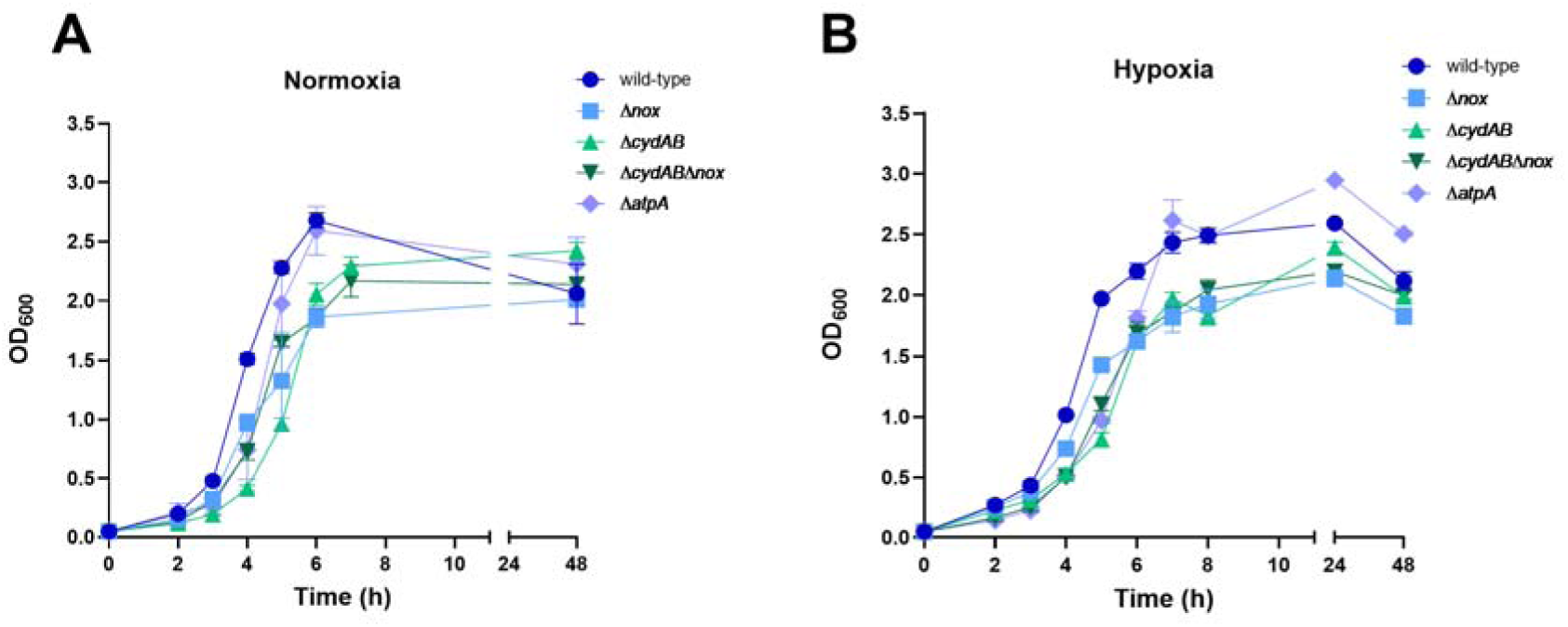
Growth of *E. faecalis* wild-type and respiratory mutants under normoxia and hypoxia. *E. faecalis* strains were grown in nutrient-rich media under normoxic (A) (aerated flask, 160 rpm, 45 % oxygen saturation) or hypoxic (B) (closed serum vials, 160 rpm, 25 % initial oxygen saturation) growth conditions. Growth was monitored as optical density over 48 h. Data is representative of the mean of biological triplicate ±SD.

To characterize the contribution of the aerobic ETC components: cytochrome *bd* oxidase and the F-type ATP synthase, as well as the cytosolic NADH oxidase in each growth condition, we first generated isogenic single gene deletion mutants of each enzyme (Δ*nox*, Δ*cydAB*, and Δ*atpA*) as well as a Δ*cydAB* Δ*nox* double deletion mutant (Table S1). All deletion mutants grew in both normoxic and hypoxic growth conditions, illustrating that no single component was essential for growth or viability under these conditions (Fig. 1).

Interestingly, there was a significant reduction in doubling time for the Δ*cydAB* and Δ*atpA* strains (T_d_ = 16.9 and 17.8 min, respectively) compared to the wild-type strain (T_d_ = 22 min) (Table S2), indicating that the ETC-deficient strains grow somewhat faster under normoxic growth conditions (Table S2).

To test if differences in growth profiles correlate with bioenergetic efficiency, we measured cell viability (CFU/mL) as a measure for fitness, cell dry weight as a measure for cell biomass (viable cells produced (g)/litre of culture), and final pH of the medium. For wild-type, the maximal biomass yield was higher in normoxic than in hypoxic cultures, suggesting that oxygen metabolism positively contributes to biomass production in *E. faecalis* (Table 1). The individual Δ*cydAB* and Δ*nox* mutants were comparable to the wild-type under both growth conditions, with similar biomass yields. However, the double deletion mutant Δ*cydAB*Δ*nox* exhibited a 3.7 – 4.3-fold reduction in stationary growth phase biomass, indicating reduced metabolic efficiency and suggesting a degree of bioenergetic redundancy between these two oxidases (Table 1). The Δ*atpA* deletion mutant also exhibited a significant reduction in biomass yield (1.6 – 2-fold), confirming that the ATP synthase contributes significantly to the cell’s bioenergetic efficiency (Table 1).

**Table 1:** Comparison of *E. faecalis* wild-type and mutants with respect to biomass yield, medium pH, and culture viability under normoxia and hypoxia growth conditions

| Strain | Biomass <sup>a</sup> |  | Culture pH <sup>b</sup> |  | CFU/mL <sup>c</sup> |  |
| --- | --- | --- | --- | --- | --- | --- |
|  | Normoxic | Hypoxic | Normoxic | Hypoxic | Normoxic | Hypoxic |
| **Wild-type** | 0.783 ± 0.10 | 0.573 ± 0.07 | 5.29 ± 0.11 | 5.50 ± 0.02 | 1.4 × 10<sup>8</sup> | 1.3 × 10<sup>8</sup> |
| **Δ*nox*** | 0.633 ± 0.17 | 0.429 ± 0.11 | 5.26 ± 0.10 | 5.60 ± 0.02 | 1.0 × 10<sup>8</sup> | 1.5 × 10<sup>8</sup> |
| **Δ*cydAB*** | 0.606 ± 0.23 | 0.524 ± 0.13 | 5.27 ± 0.04 | 5.53 ± 0.05 | 1.9 × 10<sup>8</sup> | 9.8 × 10<sup>7</sup> |
| **Δ*cydAB* Δ*nox*** | 0.453 ± 0.16* | 0.590 ± 0.07 | 5.36 ± 0.14 | 5.70 ± 0.05 | 3.2 × 10<sup>8</sup> | 3.3 × 10<sup>8</sup> |
| **Δ*atpA*** | 0.573 ± 0.04* | 0.400 ± 0.03* | 6.03 ± 0.04 | 5.73 ± 0.08 | 1.0 × 10<sup>8</sup> | 2.7 × 10<sup>8</sup> |

<sup>a</sup> Bacterial biomass yield in g of cell dry weight/liter of culture determined after 48 h of growth of three biological replicates.
<sup>b</sup> Final pH of the culture after 48 h of growth
<sup>c</sup> Bacterial cell viability determined by colony-forming units per mL after 48 h of growth.
Results are the means and standard deviations of three biological replicates and are representative of three independent experiments.
\* Statistically significant ( $p < 0.05$ ) as determined by Welch's t-test between the deletion strain and the wild-type within each defined medium condition.

### Nox is the primary oxidase responsible for oxygen consumption in *E. faecalis*

To understand the bioenergetic redundancy between CydAB and Nox we sought to understand how oxygen contributes to their bioenergetic potential. For this aim, we assayed oxygen consumption in single- and double-deletion mutants and compared it to the wild-type. First, we measured real-time oxygen concentrations in growing cultures using oxygen sensor spots (29). Briefly, sterile oxygen sensor spots were attached to the inside of the serum vial, and an optical laser was used to measure the oxygen concentration of the media at chosen time points without a need for sampling. For these experiments, we used hypoxic growth conditions to ensure consumed oxygen could not be replenished, unlike under normoxic growth conditions (Fig. S1). Crucially, a media-only control maintained consistent oxygen saturation in the absence of bacterial cells, thus verifying the experimental setup (Fig. S1).

In *E. faecalis* wild-type and Δ*cydAB* cultures, oxygen was rapidly consumed and reached extensive hypoxia after 5 h of growth (Fig. 2A and C). In contrast, we observed a significant reduction in oxygen consumption in the Δ*nox* and Δ*nox*Δ*cydAB* mutants (Fig. 2B and D), suggesting that Nox is the dominant consumer of oxygen under hypoxia- permissive conditions (Fig. 2B). However, in the Δ*cydAB* mutant culture, the oxygen levels remained somewhat higher than in the wild-type culture, suggesting a secondary role for CydAB (Fig. 2C). A minor, residual oxygen consumption was observed in both the Δ*nox* and Δ*cydAB* Δ*nox* cultures, however, *E. faecalis* encodes a heme-dependent catalase (*katA*), which may contribute to basal respiration-independent oxygen consumption (30).

**Fig. 2.**
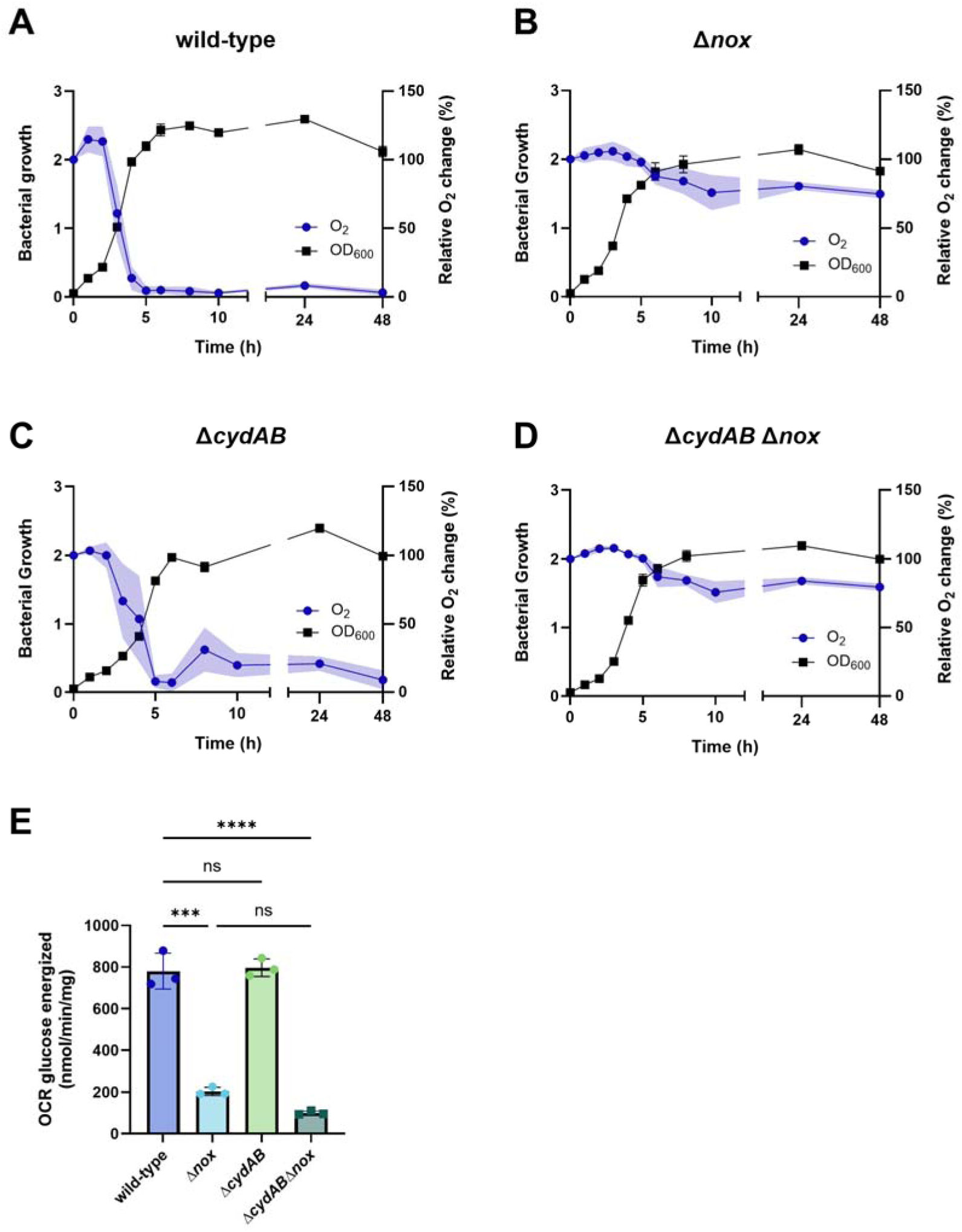
Oxygen consumption of *E. faecalis* wild-type and respiratory mutants under hypoxic growth conditions. Panels A – D: *E. faecalis* strains were grown in closed serum vials where oxygen saturation and growth were measured concurrently over time for 48 h. Graphs depict changes in relative oxygen saturation (100% = 25% oxygen saturation) alongside growth (OD_600_) for the wild-type (A), Δ*nox* (B), Δ*cydAB* (C), and Δ*cydAB*Δ*nox* (D) strains. Data is representative of the mean of biological triplicate ± SD. (E) Oxygen consumption rates (OCR, nmol of O_2_/min/mg protein) of washed cell suspension energised with glucose (10 mM) were measured for each strain. Data is representative of the mean of biological triplicate ± SD. Statistical significance was determined by an ordinary one-way ANOVA and Tukey’s multiple comparison test (p values: *** <0.005, **** <0.001, ns = not significant).

Next, we measured oxygen consumption rates (OCR) for glucose-energized, non-growing, buffered cell suspensions using the wild-type, single Δ*nox* and Δ*cydAB* mutants and Δ*cydAB*Δ*nox* double mutant. When wild-type cell suspensions were energized with glucose, the measured OCR was 780 ± 86 nmol O_2_ consumed/min/mg cell protein. Deletion of *cydAB* had no significant effect on the OCR, whereas deletion of *nox* caused a significant reduction (87%) in OCR (Fig. 2E). No significant difference was observed between the Δ*nox* and Δ*cydAB*Δ*nox* mutants (Fig. 2E). Together, these data confirm that Nox is the dominant enzyme responsible for oxygen consumption in *E. faecalis*.

To ensure that both *nox* and *cydAB* were expressed in our experimental conditions we designed gene reporter constructs of the *cydAB* (P*_cydAB_*) and *nox* (P*_nox_*) promoter regions fused to a transcriptional *lacZ* reporter and transformed these into *E. faecalis* wild-type (Table S1). Using β-galactosidase activity as a proxy for gene expression, we observed expression of both *cydAB* and *nox* under hypoxia-permissive conditions (Fig. S2).

### Semi-untargeted metabolomic analyses revealed metabolically distinct consequences of Nox and CydAB deletion

Despite distinct differences in aerobic respiration, both the Δ*nox* and Δ*cydAB* mutants showed wild-type-like growth and biomass production. Therefore, to determine the global metabolic role of oxygen consumption facilitated by Nox and CydAB, semi-untargeted metabolomics was carried out on stationary-phase normoxia-adapted (45 % oxygen saturation) and hypoxia-adapted (>1 % oxygen saturation) wild-type, Δ*nox* and Δ*cydAB* cultures. Metabolomic analysis of *E. faecalis* wild-type adaptation to normoxia and hypoxia showed a clear distinction between these conditions indicative of oxygen concentration-dependent metabolic states, consistent with a previous report by Portela *et al.* (Fig. S3 and Table S4) (31).

Hierarchical clustering of the wild-type, Δ*nox* and Δ*cydAB* strains showed that the metabolic profile of each mutant was distinct from the wild-type, and independent of growth conditions. This highlights that deletion of *nox* or *cydAB* has a greater impact on the metabolic profile than oxygen tension itself (Fig. 3A and S4). The Δ*nox* strain exhibited an increase in the NADH:NAD^+^ ratio relative to the wild-type, consistent with its annotated role in regenerating NAD^+^ (Fig. 3B). The Δ*nox* also exhibited a reciprocal decrease in ATP:ADP ratios (Fig. 3C). Further analysis revealed an accumulation of upper glycolytic intermediates, including glucose-6-phosphate and fructose-6-phosphate, and reciprocal depletion of downstream glycolytic metabolites, such as 3-phosphoglycerate and phosphoenolpyruvate (Fig. 4 and Table S4). These metabolites indicate a distinct division from the NAD^+^-dependent enzyme glyceraldehyde-3-phosphate dehydrogenase in the Δ*nox* strain. This suggests that loss of Nox compromises cytoplasmic NADH oxidation, resulting in a reduction of free NAD^+^ that restricts glycolytic flux and associated ATP production. Accumulations of the TCA cycle metabolites: cis-aconitate and itaconate (in the absence of a corresponding increase in succinate), and the reduced derivative of α- ketoglutarate, 2-hydroxyglutarate are similarly indicative of NAD^+^-limited decreases in activity of the NAD^+^-dependent enzyme, isocitrate dehydrogenase (Fig. 4 and Table S4).

**Fig 3.**
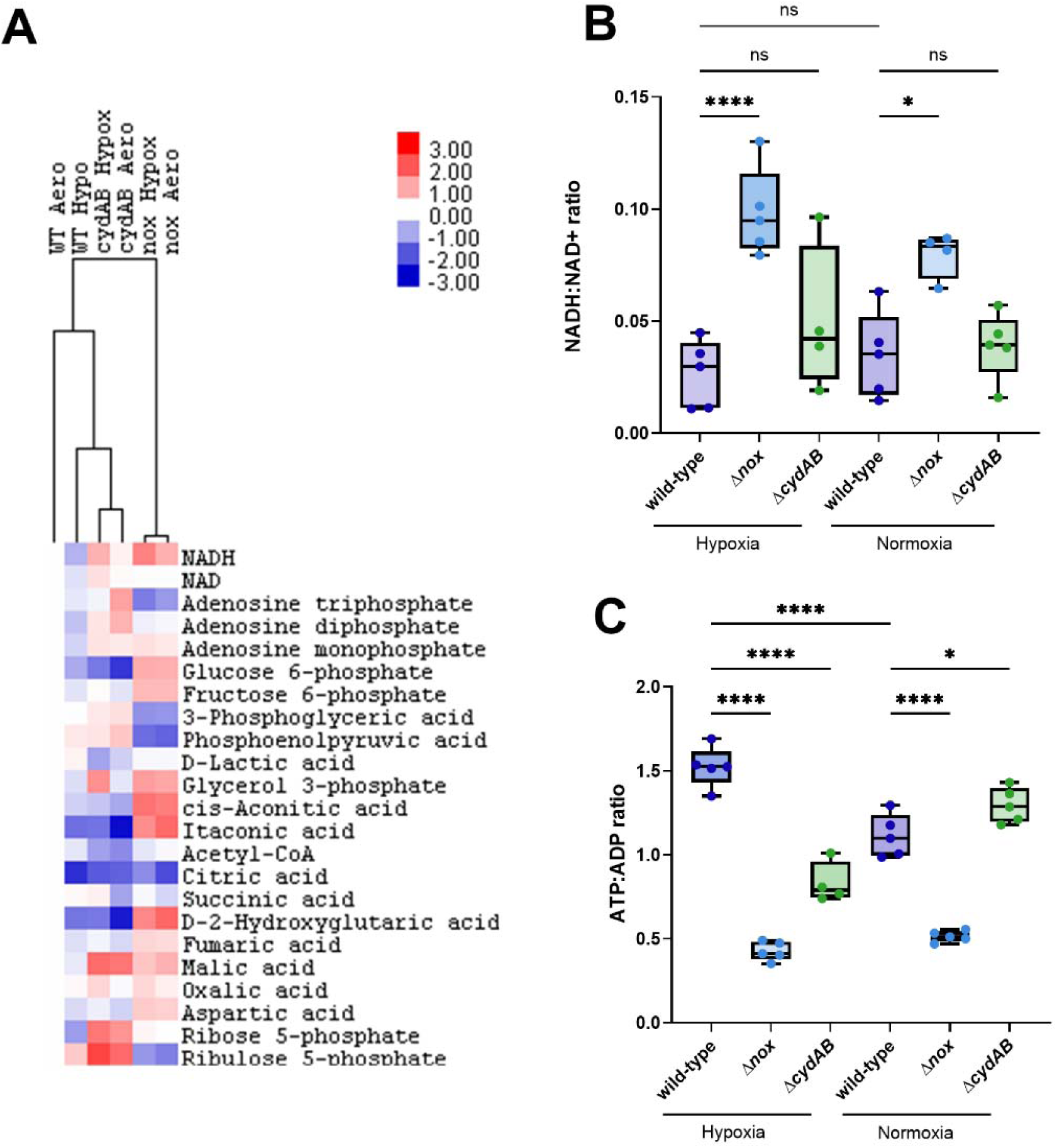
Semi-targeted metabolomics of *E. faecalis* wild-type and mutants Δ*nox* and Δ*cydAB* grown in normoxic and hypoxic conditions. *E. faecalis* wild-type (WT), Δ*nox* (nox) and Δ*cydAB* (cydAB) strains were grown to stationary phase (9 h) to reach normoxic (Aero) and hypoxic-adapted (Hypo) conditions. Semi-targeted metabolomics was carried out on intracellular metabolites for each strain and condition. The mean metabolic changes for selected metabolites have been clustered based on fold-change using Metabolanalyst (A). Data is representative of the mean of five biological replicates. Ratios of NADH/NAD^+^ (B) and ATP/ADP (C) have been plotted for each strain and growth condition. Data is representative of the median of five biological replicates. Statistical significance was determined using a one-way ANOVA and Tukey’s multiple comparison test (p values: * <0.05, **** <0.0001, ns = not significant).

**Fig. 4.**
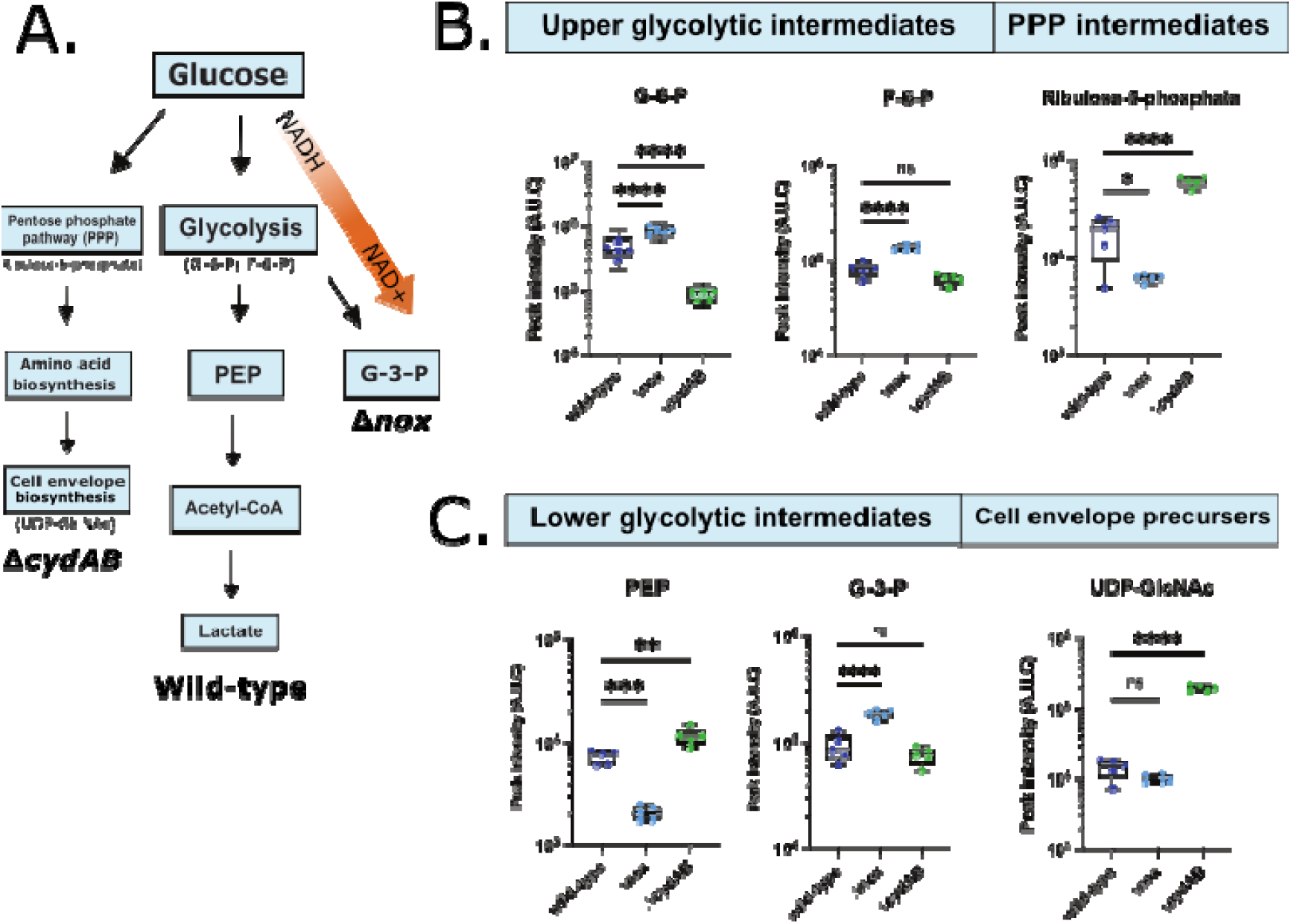
Key pathway changes of *E. faecalis* wild-type and mutants Δ*nox* and Δ*cydAB*. Panel A represents a simplified pathway of central carbon metabolism and the consequences for each oxidase deletion from pathway enrichment analysis of semi-targeted metabolomics. Panel B shows the initial upstream rerouting from glucose to either glycolysis or the pentose phosphate pathway. Panel C shows the metabolic consequences downstream of glycolysis or of the pentose phosphate pathway. Samples used 5 biological replicates, and statistical significance was determined using (A-B) a Student’s t-test or (C-D and F-G) a one-way ANOVA with Tukey’s multiple comparison test, where ** denotes p<0.01, *** denotes p<0.005, **** denotes p<0.001, and ns indicates not significant. Acronyms used in this figure: G-6-P (glucose-6-phosphate), F-6-P (fructose-6-phosphate), PEP (phosphoenolpyruvate), G-3-P (glycerol-3-phosphate) and UDP-GlcNAc (Uridine diphosphate N-acetylglucosamine).

These metabolic changes suggest that the intracellular redox state cannot be relieved as glycolytic carbon flow through the incomplete TCA cycle is restricted. These metabolic changes indicate that Nox-mediated regeneration of cellular NAD^+^ pools is required to sustain glycolytic carbon flow and oxidative phosphorylation.

In contrast, the Δ*cydAB* deletion strain exhibited a metabolic profile similar to that of wild-type adapted to hypoxia with comparable ATP:ADP and NADH:NAD^+^ ratios, indicative of an overall preservation of energetic and redox homeostasis (Fig. 3). The Δ*cydAB* deletion exhibited increased levels of pentose phosphate pathway intermediates, including ribulose-5-phosphate and ribose-5-phosphate (Fig. 4). This partial redistribution of carbon from glycolysis towards the pentose phosphate pathway may help adapt to a more restricted respiratory capacity. These findings, thus, demonstrate that Nox is the dominant oxidase responsible for maintaining NAD⁺ regeneration and central metabolic flux in *E. faecalis*, whereas CydAB contributes to a distinct, oxygen-responsive metabolic program that is not essential for bulk redox or energetic homeostasis under these conditions.

### The F-type ATP synthase functions primarily as an ATP-driven proton pump to generate proton motive force in *E. faecalis*

We have demonstrated that Nox and Cyd contribute to bioenergetic efficiency and carbon metabolism in *E. faecalis*. Next, we wanted to investigate whether these components, along with the ATP synthase, play a role in the key bioenergetic process of generating PMF. The PMF is generally assumed to play an essential biological role in growing and non-growing cells (32–37). However, it is currently unknown how the different respiratory components of *E. faecalis* contribute to the generation of PMF, or its constituent parts, the transmembrane electric potential (ΔΨ) and the transmembrane pH difference (ΔpH).

The voltage-sensitive fluorescent probe DiSC_3_(5) detects the ΔΨ at a single-cell level (38,39). First, we determined that DiSC_3_(5) fluorescence can be used to monitor membrane potential in *E. faecalis* wild-type, building on protocols previously established in other bacterial species (Fig. 5A and B). While we observed fluorescence, this was markedly lower than those observed in other Gram-positive species such as *Staphylococcus aureus* (Fig. S5). In principle, this could be due to low permeability of DiSC_3_(5) across the enterococcal cell envelope, or it could also indicate that *E. faecalis* possess an innately low ΔΨ, as previously suggested (40,41). To investigate this, we challenged *E. faecalis* wild-type with the membrane depolarising ionophore gramicidin and the ΔpH-dissipating ionophore nigericin. Gramicidin is an antimicrobial peptide that generates small cation-specific (predominantly K^+^ and H^+^) membrane channels that lead to the collapse of ΔΨ. Incubation with gramicidin resulted in membrane depolarisation, which confirmed voltage-dependent staining in *E. faecalis* by DiSC_3_(5) (Fig. 5A). Nigericin, in contrast, mediates an electroneutral exchange of K^+^ and H^+^ ions, resulting in a dissipation of the pH gradient (ΔpH) (42). If *E. faecalis* maintains an innately low membrane potential due to a high ΔpH, an addition of nigericin should allow cells to generate a higher ΔΨ in response to the dissipated ΔpH. Incubation with nigericin triggered an increase in membrane potential, as indicated by high DiSC3(5) fluorescence signals (Fig. 5A and B, Fig. S6). These findings indicate that *E. faecalis* indeed maintains a high ΔpH, which acts as the dominant component of PMF.

**Fig. 5.**
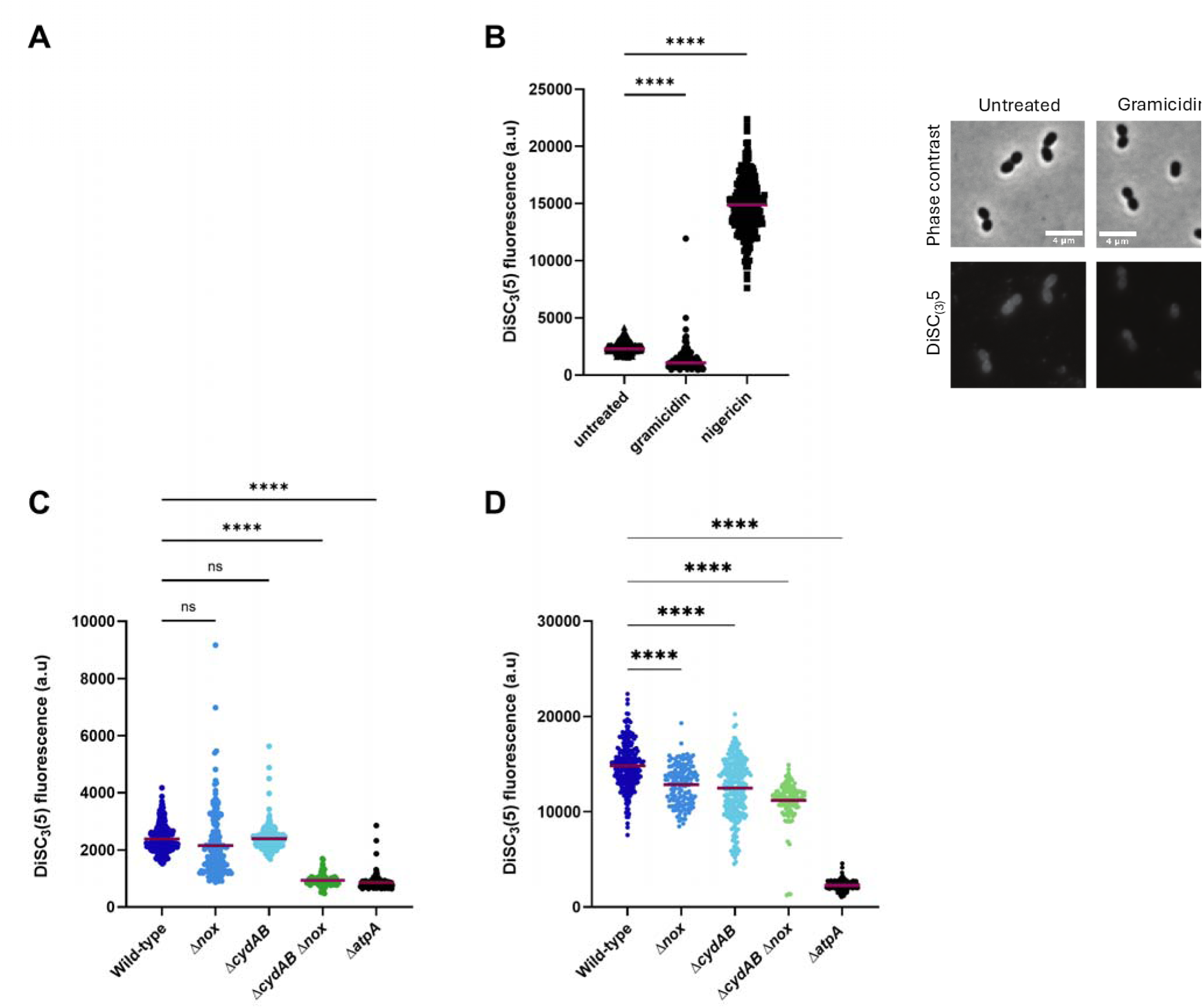
Single-cell measurements of membrane potential for wild-type and respiratory mutants. *E. faecalis* strains were grown to early-mid-log phase under normoxic growth conditions and incubated with the voltage-sensitive dye DiSC_3_(5). Cell membrane potential levels (ΔΨ) were determined in the wild-type by single-cell fluorescence microscopy (A and B), with the membrane-depolarising ionophore gramicidin (10 µM) and ΔpH-dissipating ionophore nigericin (10 µM) acting as controls. Single-cell membrane potential levels were determined for strains deficient in NADH oxidase (Δ*nox*), cytochrome *bd* oxidase (Δ*cydAB*), and F-type ATPase (Δ*atpA*), and compared to wild-type cells (C). Cells were treated with nigericin (10 µM) to determine hyperpolarisation upon ΔpH collapse for each strain (D). Data is plotted as the median (red line) with individual data points. Statistical significance was determined by one-way ANOVA with Tukey’s multiple comparison. P value: **** <0.0001 and ns = not significant.

To determine whether the respiratory components contribute to PMF, DiSC_3_(5) microscopy was carried out on each of the deletion mutants Δ*nox*, Δ*cydAB*, Δ*atpA* and the double deletion mutant Δ*cydAB* Δ*nox*. While ΔΨ was significantly decreased in the Δ*atpA* and the Δ*cydAB* Δ*nox* strains, the absence of Nox or CydAB individually had no effect on ΔΨ (Fig. 5C). These data suggest that the ATP synthase plays a dominant role in generating ΔΨ, with the metabolic disturbances observed in Δ*cydAB* Δ*nox* likely indirectly hampering the ability of ATP synthase to maintain ΔΨ. To further characterise how the different components contribute to overall PMF, each deletion strain was incubated with nigericin followed by DiSC_3_(5) microscopy. Here, all strains showed strong nigericin-dependent hyper-polarisation except the Δ*atpA* deletion mutant (Fig. 5D), suggesting that high ΔpH is maintained in all strains except Δ*atpA.* In summary, these results suggest that while Nox and CydAB collectively contribute to PMF generation, they likely do so indirectly via the F-type ATPase, the main component of *E. faecalis* responsible for generating both ΔΨ and ΔpH components of the PMF through ATP-driven proton transport.

## Discussion

*E. faecalis* is a highly successful generalist bacterium which grows as a commensal in the human and animal gut (4,43). However, it is also one of the leading causes of several serious hospital-acquired infections (44,45). Targeting metabolism has been increasingly used to great benefit in key pathogens including *Mycobacterium tuberculosis,* as seen with the introduction of ATP synthase inhibitors in first-line drug treatment (46,47). Despite the widespread importance of aerobic respiration in bacterial physiology, including in well-characterised pathogens such as *S. aureus*, *Listeria monocytogenes*, and *M. tuberculosis*, its role in the facultative anaerobe *E. faecalis* remains poorly defined (48–51).

Here, we characterise the bioenergetic roles of the aerobic respiratory enzymes Nox, CydAB and F-type ATPase in *E. faecalis*. To do this, we systematically generated isogenic genetic mutants of each enzyme and measured growth parameters to conclusively show that their activity is not critical to overall cell growth. Furthermore, the CydAB- and F-type ATPase-deficient strains displayed a quicker exponential doubling time, suggesting that a functional ETC is a rate-limiting factor under aerobic growth conditions in *E. faecalis.* This contrasts with classical models of bacterial respiration, in which oxidative phosphorylation enhances growth efficiency through increased ATP yield (22,52). Instead, our findings support a model in which *E. faecalis* prioritizes rapid fermentative metabolism over respiratory efficiency, particularly in nutrient-rich environments (53).

Enterococci are commonly found in the gastrointestinal tract, which is a highly competitive, hypoxic environment. Recent work has demonstrated that during *Clostridioides difficile* infection, *E. faecalis* acquires host heme by toxin-mediated tissue damage and uses it to reconstitute the CydAB which confers a fitness *in vivo* (54). Similarly, *E. faecalis* encodes an anaerobic heme-degrading enzyme (anaerobilin synthase) that mediates iron release from heme under anaerobic conditions, which has been demonstrated to aid gut colonization and virulence in multiple infection models (55). Collectively, these findings indicate that the primary role of the respiratory chain might instead be to provide a competitive fitness advantage within heterogeneous host environments, where nutrient limitation, immune defences, and dynamic oxygen availability create ecological niches that *E. faecalis* is adapted to exploit. This may explain why we do not observe significant differences in the growth phenotypes tested under laboratory conditions. Testing these mutants in a polymicrobial environment could prove interesting in the future.

Although *E. faecalis* is a facultative anaerobic bacterium, it contains two oxidase enzymes (Nox and CydAB) (10,27). Here we show that Nox is the primary oxidase in *E. faecalis* responsible for bulk oxygen consumption, and that the loss of respiratory capacity under high oxygen tension does not significantly affect growth. We therefore propose that *E. faecalis* primarily operates an anaerobic metabolic state and utilizes Nox to rapidly sequester oxygen and sustains a high NAD^+^ pool for anaerobic metabolism. The metabolomics data suggest that this oxygen-adaptation strategy supports fermentative metabolism, redox balancing, and oxidative stress defence in both anaerobic and aerobic environments. Indeed, previous continuous culture metabolomics and ¹³C-labelling work demonstrated that rapid transition of *E. faecalis* from anaerobic to aerobic conditions upregulated glycolysis approximately two-fold (31).

Unlike close relatives such as *E. faecium*, *E. faecalis* maintains the core components of an active ETC (10). The PMF is generally assumed to play an essential biological role in growing and non-growing cells (32–37). Therefore, we sought to determine whether the primary contribution of ETC upon aerobic respiration was to generate PMF (10,35,56). In doing so, we first demonstrated that *E. faecalis* maintains a remarkably low steady-state membrane potential. We further showed that both the Δ*atpA* deletion and double oxidase deletion strains exhibited an even further reduced ΔΨ, with the Δ*atpA* mutant exhibiting a reduced ΔpH as well. These results thus provide evidence that the *E. faecalis* F-type ATPase may, in fact, act as an ATP-driven proton pump that serves as the cell’s primary mechanism to generate PMF. Which is a highly atypically mechanism and does so alongside active aerobic respiration (Fig. 6). While a PMF-generating role has been previously reported for the F-type ATPase in *E. hirae*, this species does not encode an ETC and thus grows fermentatively (17,57,58). Work in *E. hirae* has shown that its V-type ATPase can synthesise ATP when driven by a sodium motive force, suggesting that enterococci may have greater versatility in ion homeostasis and associated bioenergetic processes than previously assumed (59). That said, it is all the more unexpected and remarkable that loss of the F-type ATPase is not detrimental to *E. faecalis*, perhaps indicative of a bioenergetically robust but parsimonious physiology. Rather than being involved in PMF generation, our data argue that the primary metabolic role of the *E. faecalis* ETC lies in facilitating re-oxidation of NADH and, somewhat paradoxically, optimizing glycolytic metabolism.

**Fig. 6.**
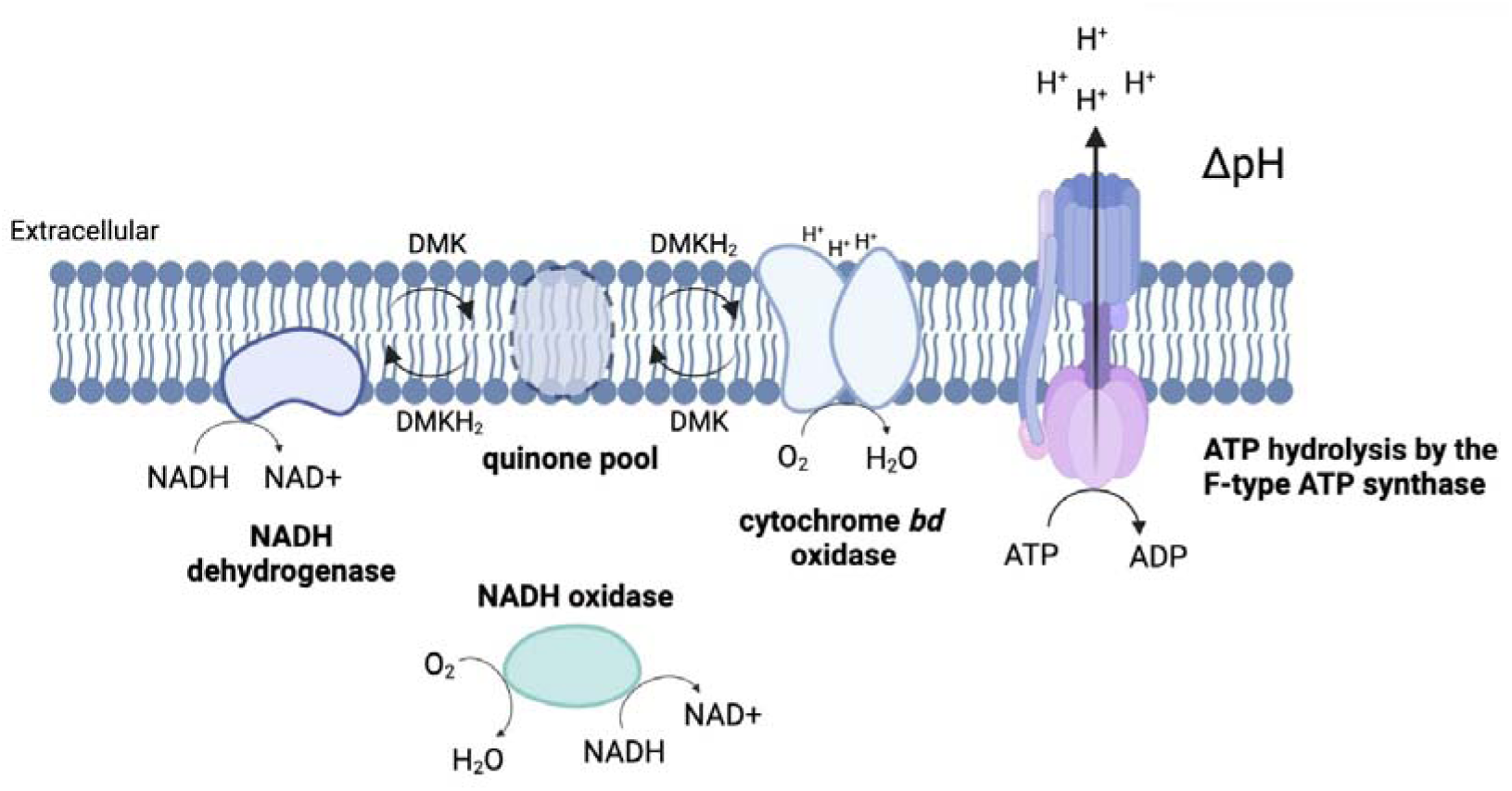
Schematic of the respiratory complexes in *E. faecalis*. The cytosolic NADH oxidase (*nox*; EF1586) is a major consumer of oxygen and plays a significant role in NADH turnover in *E. faecalis*. The terminal cytochrome *bd* oxidase (*cydAB*; EF2060-2061) cycles demethylmenaquinone (DMK/DMKH_2_) powered by the reducing equivalent of NADH through the type II NADH dehydrogenase (*ndh2*; EF2055). We propose that the F-type ATP synthase (*atpA-H*; EF2614-2607) acts to hydrolyse rather than synthesise ATP to actively drive proton export, allowing proton motive force to be maintained during fermentation. This is evidenced by hyper-polarisation of the cell upon addition of the ΔpH-dissipating ionophore nigericin.

*E. faecalis* is a highly adaptable generalist bacterium that occupies both commensal and pathogenic niches. Its success likely stems in part from this metabolic flexibility, enabling survival across fluctuating oxygen and nutrient conditions. Our findings challenge the assumption that aerobic respiration universally confers a growth advantage and instead highlight the central importance of redox balance and particularly the NADH:NAD⁺ ratio in shaping metabolic flux. Perhaps for *E. faecalis* and other competitive facultative anaerobic bacteria, the preservation of these redox redundancies provides resilience to a broader niche range including pH, osmotic pressure, and host derived immune oxidative stress. By defining the limited but distinct roles of Nox, CydAB, and the F-type ATPase, this work provides a foundation for future efforts to exploit metabolic vulnerabilities in *E. faecalis*, particularly given that key antimicrobials such as daptomycin and the human host defense peptide LL-37, which target membrane energetics (60).

In summary, we have shown that *E. faecalis* makes only limited use of the aerobic respiratory chain for energy generation and instead propose that maintaining a favourable NADH:NAD^+^ ratio linked to oxygen consumption plays a much greater role in dictating this bacterium’s bioenergetic strategy. Moreover, we have shown that *E. faecalis* maintains PMF with a dominant contribution of ΔpH in an ATP-dependent manner through the F-type ATPase even under aerobic respiration and highlighted flexible bioenergetic networks that favour metabolic flexibility rather than maximal ATP yield. Together, our findings support a revised model in which intracellular redox balance, rather than oxidative phosphorylation, is the principal determinant of bioenergetic strategy in *E. faecalis*.

### Materials and Methods Growth of *E. faecalis* strains

The strains and plasmids used to generate the knockout constructs were shown (Table S1). *E. faecalis* strains were grown in rich Brain Heart Infusion (BHI) medium. All *E. faecalis* agar plate experiments were carried out on BHI agar unless otherwise stated. *E. coli* strains were grown in Luria-Bertani (LB) broth and plated on LB agar unless otherwise stated. All antibiotic stocks were made up to 10 mg/mL stock in MilliQ-dH_2_O (Sigma Aldrich). Nigericin was prepared in 100% absolute grade ethanol to 10 mM and used at a final working concentration of 10 µM in 1% ethanol. The voltage-sensitive membrane-permeable dye DiSC_3_(5) was made up in DMSO at 10 mM for a final working concentration of 1 µM and 1% DMSO.

### Construction of *E. faecalis* mutants

All *E. faecalis* knockout strains were generated using pIMAYZ plasmid, and chloramphenicol (10 µg/mL final concentration) was added to the media to select for this plasmid (61). This was carried out as previously described (61,62). In summary, to generate all *E. faecalis* mutants, homologous end recombination was used with the temperature-sensitive pIMAYZ knockout plasmid. Flanking regions up- and downstream of each gene of interest (GOI) were amplified using primers shown (Table S2). The GOI flanks were ligated into the pIMAYZ plasmid, then amplified in *E. coli* to allow plasmid amplification. The pIMAYZ_GOI knockout plasmid was transformed into electrocompetent *E. faecalis* JH2-2, and the recombination event screened using IM151 and IM152 primers shown (Table S2). The double oxidase mutant (Δ*cydAB* Δ*nox*) was generated in a Δ*cydAB* background, using electrocompetent Δ*cydAB* transformed with the pIMAYZ_nox plasmid. All deletion mutants generated in this study were confirmed to be clean deletions with no additional single nucleotide polymorphisms or genetic rearrangements to account for any phenotypes.

### Growth phenotypes measured

Dry weight was taken as the total volume of bacterial broth washed and concentrated down to 2 mL before being transferred to tin foil weight boats and left to dry for 72 h at 100 °C. A media-only control was used to find the standard weight after dehydration. Tin foil weight boats were individually weighed prior to the addition of culture to give the initial weight (g_i_). Weight was measured rapidly after dehydration to measure the final dry weight (g_f_), and the following formula was used to determine the final cell biomass. Cell colony-forming units (CFU) and optical density (OD_600_) measurements were taken using standard practice. Culture pH was determined after 48 h growth.

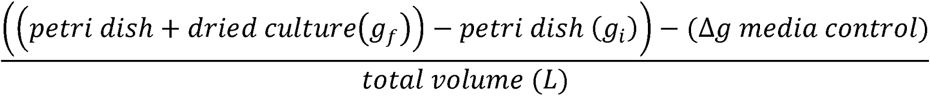

Extracts were taken during exponential phase to calculate the specific exponential growth rate (µ), which was then used to calculate the doubling time (Td), as shown in the formulas below.

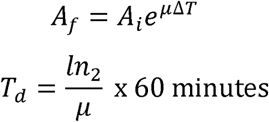

### Oxygen consumption measures

Initial oxygen consumption measurements were carried out in washed whole-cell suspensions using a fluorespirometer (OROBOROS Oxygraph-O2K, Innsbruck, Austria) in a sealed chamber containing a Clark-type oxygen electrode. These washed *E. faecalis* cells were resuspended in 1× PBS to an OD_600_ of 0.5 to de-energise the cells. Before each experiment, cultures were equilibrated with ambient air and spiked with a carbon source (10 mM glucose f/c) to re-energise the cells. All measurements were taken at 37 °C, 750 rpm, and data was recorded at 1 s intervals. Once oxygen consumption reached a stable rate, the average oxygen consumption was measured. BSA assays were used to normalize protein concentrations in the cell suspensions to determine the rate of OCR per mg of protein.

Oxygen consumption in serum vials and flasks was measured using the stand-alone fibre optic trace oxygen meter Fibox 4 trace coupled with the oxygen sensor spots pst7-10YAU- P5-YP (PreSens Precision sensing GmbH Am Biopark, Germany). These sensors were chosen as they covered a 100 %-0.1 % oxygen range and were autoclavable. The sensors were glued at the bottom of serum vials and flasks. Cultures were inoculated into fresh media to an approximate OD_600_ of 0.05, allowing 2/3 headspace. Cultures were grown in biological triplicate (37 °C with 160 rpm) in rubber-stopped serum vials (hypoxia) or flasks (normoxia). Oxygen concentration was measured every h until stationary phase was reached. Growth was measured in parallel with samples taken from the serum vials using a hypodermic needle that was inserted through the sterilized rubber stopper.

### **β**-galactosidase assays

The upstream region of each GOI was selected starting before the ATG start codon and the sequence for EcoRI and BamHI restriction cut sites were added. These were then ordered with GeneScript using their custom vector construction to synthesize the pTCVlac- GOI constructs. These constructs were then transformed into electrocompetent *E. faecalis* wild-type cells.

To quantitatively determine the level of GOI induction under specific growth conditions, *E. faecalis* wild-type cells were grown in serum vials with oxygen monitoring (sensor-spots). Extracts were taken from these serum vials at 2, 4, 6, 8, and 10 h with OD_600_ taken to normalize cells for the β-galactosidase assay using the formula below. At each time point, extracts were harvested by centrifugation and stored at -20 °C. β-galactosidase assays were carried out as previously described and gene expression determined using the formula below (63).

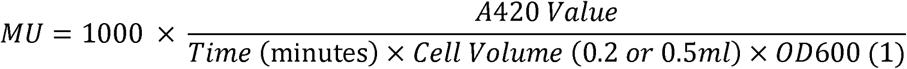

### Membrane potential fluorescence microscopy

For standard single-measure microscopy, cells were grown to mid-exponential phase in Mueller-Hinton or Brain Heart Infusion at 37 °C at 200 rpm. Once logarithmic growth phase was reached, 100 µL of the growing culture was transferred to 2 mL Eppendorfs containing a perforated lid to support aeration and grown with 600 rpm of aeration at 37 °C on a bench-top thermomixer. The cells were then incubated with the voltage-sensitive dye DiSC_3_(5) for 5 min at a final concentration of 1 µM following previously established guidelines (38). Fluorescence microscopy sample preparation was carried out briefly as follows: a 1.2 % agarose: dH_2_O suspension was pipetted onto Teflon-coated multi-spot microscope slides and warmed to room temperature prior to use. The DiSC_3_(5)-treated cell suspension (0.5 µL) was pipetted onto an agarose surface and evaporated at 37 °C (∼2 min) before adding a coverslip. Oxygen loss under coverslips has previously been reported (64). To minimise this effect, images were taken within 10 min of coverslip addition. Microscopy images were taken with a Nikon Eclipse Ti (Nikon plan Apo 100 x containing 1.40 Oil Ph3 objective lens and CoolLED pE-4000 light source). The excitation/emission filter used for DiSC(_3_)5 microscopy was the Cy5 filter, which excites at 628 nm ±40 and measures emission at 692 nm ±40, with 100 % power of a 635 nm LED line.

For all fluorescence microscopy, images were analysed using ImageJ/Fiji. A semi-automated pipeline was written and optimised for the use of *E. faecalis* to quantify the fluorescence. This pipeline was adapted from the pre-established *B. subtilis* image analysis code used from GitHub https://doi.org/10.5281/zenodo.11388986. The individual cells were selected as regions of interest (ROIs), Watershed and fill hole functions of ImageJ were used to individually identify each bacterium using the corresponding phase contrast images. The background fluorescence signal was then removed (using average fluorescence outside ROIs) to set the threshold for the fluorescence channel, and these ROIs were then measured for their own individual mean fluorescence values.

### Intracellular metabolite extraction

The sampling, quenching and intracellular metabolic extraction was carried out using previously published guidelines (65). All extracts were harvested after 9 h, using oxygen-sensor spots to confirm that oxygen saturation in the media was 1 % (hypoxic - adapted) or >40 % (normoxia - adapted). In summary, 40 mL of culture in five biological replicates was centrifuged down at 4 °C and then resuspended with ice-cold glycerol saline solution (3:2) to quench metabolic activity. Samples were then pelleted by centrifugation at -20 °C. Samples were washed in cold 1:1 glycerol saline in triplicate, resuspended in 2 mL dH2O, and decanted into fresh (glycerol-free) falcons. The samples were washed a final time with dH_2_O before normalized to OD_600_ 5 and OD_600_ 0.5 in 4:1 methanol:dH_2_O. Following resuspension, an internal CHAPS-HEPES-TRIS-PIPES (2 µM) standard was added to each sample and the extraction buffer was sent as a negative control to Metabolomics Australia. Intracellular metabolites were then extracted using three freeze-thaw cycles.

### Semi-untargeted metabolomics

Samples were run in a biologically untargeted fashion using an in-house list of 550 potential metabolites, which provided known peaks alongside our unknown samples, by Metabolomics Australia. The chromatography conditions were modified from Lu *et al.* (66). Metabolites were separated and detected using a Vanquish Horizon UHPLC system (Thermo Scientific) and run on a Orbitrap-ID-X Tribrid mass spectrometer (Thermo Scientific). Briefly, separation was performed using a SeQuant *zic*-pHILIC column (150 mm × 4.6 mm, 5 µm particle size; Merck) at 25 °C, with a binary gradient of solvent A (20 mM ammonium carbonate (pH 9.0; Sigma-Aldrich) and solvent B (100% acetonitrile (Merck, # 100029). A gradient of A/B solvents was run at a flow rate of 300 μL/min as follows: 0.0 min, 80 % B; 0.5 min, 80 % B; 15.5 min, 50 % B; 17.5 min, 30 % B; 18.5 min, 5 %; 21 min, 5 % B; 23–33 min, 80 % A.

For metabolite detection, the Orbitrap ID-X Tribrid Mass Spectrometer (Thermo Scientific) was coupled to a heated electrospray ionization source and performed as follows: sheath gas flow 40 arbitrary units, auxiliary gas flow 10 arbitrary units, sweep gas flow 1 arbitrary units, ion transfer tube temperature 275 °C, and vaporizer temperature 320 °C. The radio frequency lens value was 35 %. Data was acquired in negative polarity with spray voltages of 3500 V. Samples were run in a random order, and the quality of data produced was assessed by the peak variation of pooled samples (all samples combined equally) every 5 samples. Batch-to-batch variation was controlled using routine blank runs, and media blank runs between each sample batch and the internal standard was run during every study sample. The data was collected using Thermo Tracefinder (V 4.1) (General Quan Browser). Metabolites were assigned to sample peaks in El-Maven v.0.12.1 by comparison to the peaks in the standard library (67).

Identified metabolites of all samples were provided as raw values as peak intensity determined by area under the curve. Raw data were processed and analysed using the MetaboAnalyst v6.0 web server (https://www.metaboanalyst.ca/docs/About.xhtml) (68). The internal standard of CHAPS-TRIS-MOPS-PIPES were run alongside the analysis of each intracellular metabolome to ensure comparable extraction. Peak intensities of all identified metabolites in a combined dataset of all samples were processed using the Statistical Analysis [one factor] pipeline. Here, the dataset was median normalized across all experimental conditions and strains and log_10_ transformed. Key comparisons shown are between the wild-type normoxia and wild-type hypoxia adapted growth conditions of metabolites with fold changes >1.5 × (p<0.1) shown as relative abundance. Comparisons between all strains in normoxia conditions were compared and metabolites with fold changes >1.5 × (p<0.1) shown as relative abundance. Hierarchically clustered heatmaps of the processed metabolomics data were generated using Java TreeView.

## Declarations

We declare no conflicts of interest.

## Acknowledgements

We are grateful for the following people for their discussions and academic input: Chen-ye Cheung, Will Jowsey, and David Mayo Muñoz. We would also like to acknowledge the support from the New Zealand Health Research Council, the Royal Society Catalyst Grant, Otago PhD scholarship and the Elman-Poole Traveling Scholarship for their funding support.

## Supplementary Tables

**Table S1:** List of bacterial strains and plasmids used in this study.

| Strain | Genetic background | Reference |
| --- | --- | --- |
| *E. faecalis* |  |  |
| JH2-2 | Wild-type *E. faecalis* strain with Rif<sup>R</sup> Fus<sup>R</sup> | (1) |
| $\Delta nox$ | Deletion of *nox* (EF1586) in JH2-2 | This study |
| $\Delta cydAB$ | Deletion of *cydAB* (EF2060-EF2061) in JH2-2 | This study |
| $\Delta cydAB\Delta nox$ | Deletion of *nox* in a $\Delta cydAB$ genetic background | This study |
| $\Delta atpA$ | Deletion of alpha subunit of F-type ATP synthase (EF2610) | This study |
| *E. coli* |  |  |
| DH10B | *E. coli* strain used to propagate pIMAYZ plasmid | (2) |
| Plasmids | Selective markers | Reference |
| pIMAYZ | Cm<sup>R</sup> | (3) |
| pTCVlac | Erm<sup>R</sup> | (4) |

**Table S2:** Maximum specific doubling time (T_d_ = min) of *E. faecalis* versus mutants deficient in respiratory enzymes in normoxic growth conditions (45% oxygenation shaken flasks). Data is representative of the mean ± SD of biological triplicate.

| | Wild-type | $\Delta nox$ | $\Delta cydAB$ | $\Delta cydAB \Delta nox$ | $\Delta atpA$ |
| --- | --- | --- | --- | --- | --- |
| Normoxia | 22.4 ± 2.2 | 21.6 ± 0.8 | 16.9 ± 0.9* | 20.6 ± 1.0 | 17.8 ± 2.2* |

\* Statistically significant ( $p < 0.01$ ) as determined by one-way ANOVA.

**Table S3:**
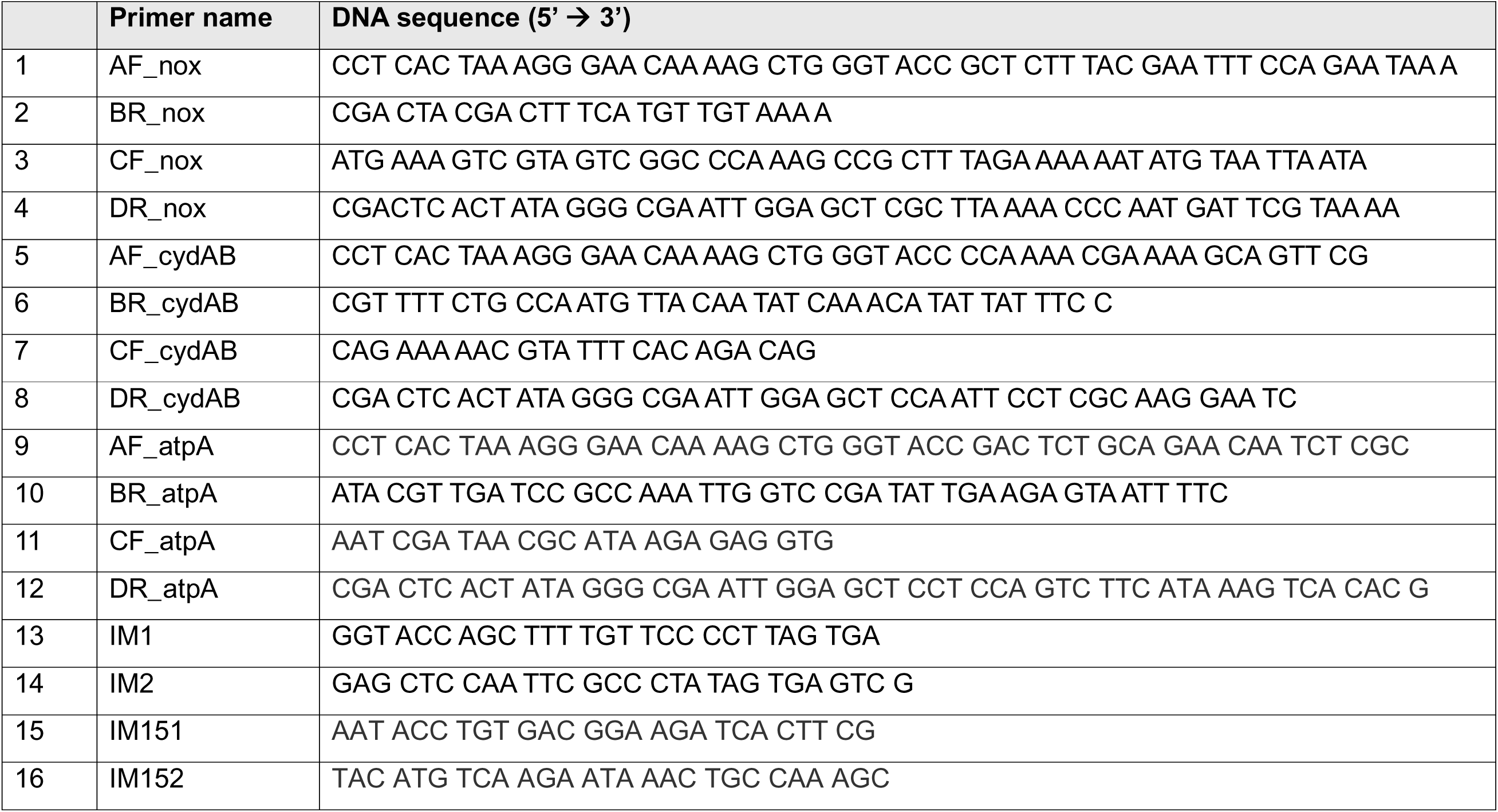
List of primers used in this study.

## Supplementary Figures

**Fig. S1.**
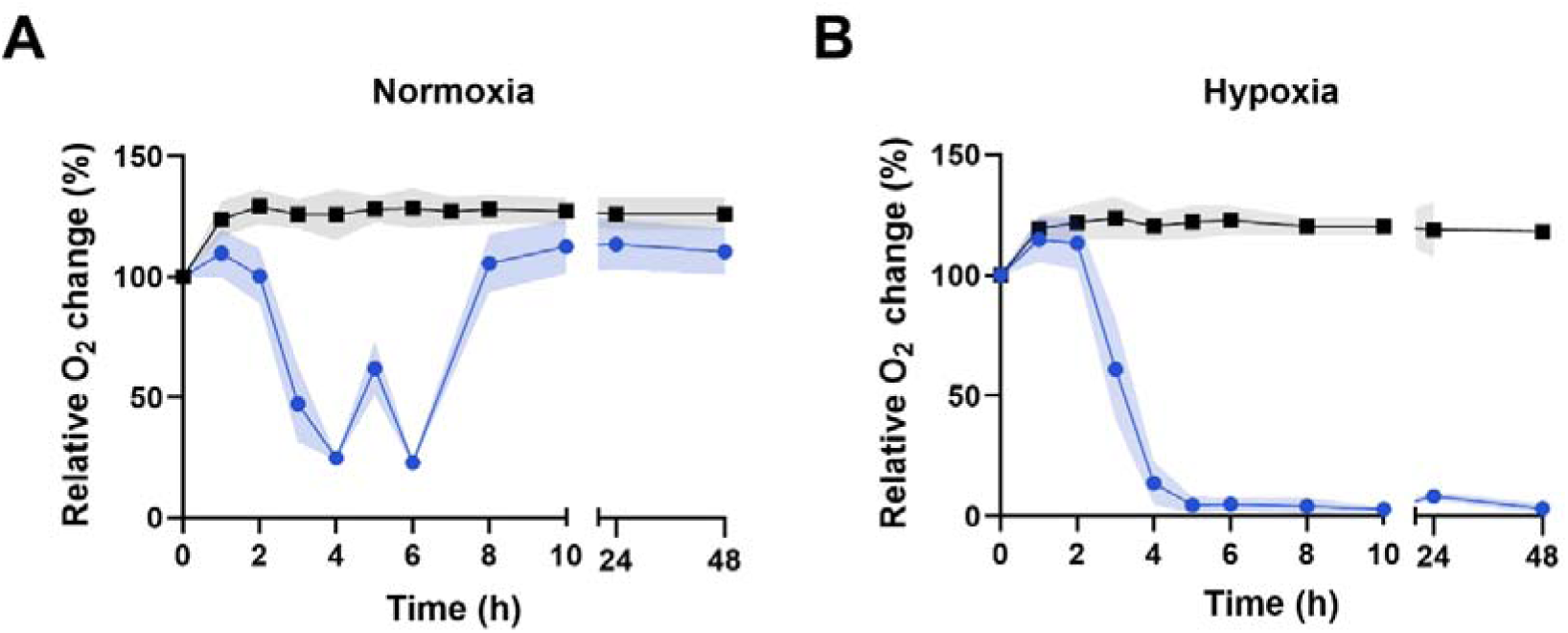
Oxygen consumption of *E. faecalis* wild-type under normoxic and hypoxic growth conditions. Nutrient-rich BHI medium was inoculated with (blue dots) and without (black squares) wild-type *E. faecalis*. Relative oxygen saturation was measured under (A) normoxia (100% = 45%) and (B) hypoxia (100% = 25%) over a period of 48 h. Data is representative of the mean of biological triplicate ± SD.

**Fig. S2.**
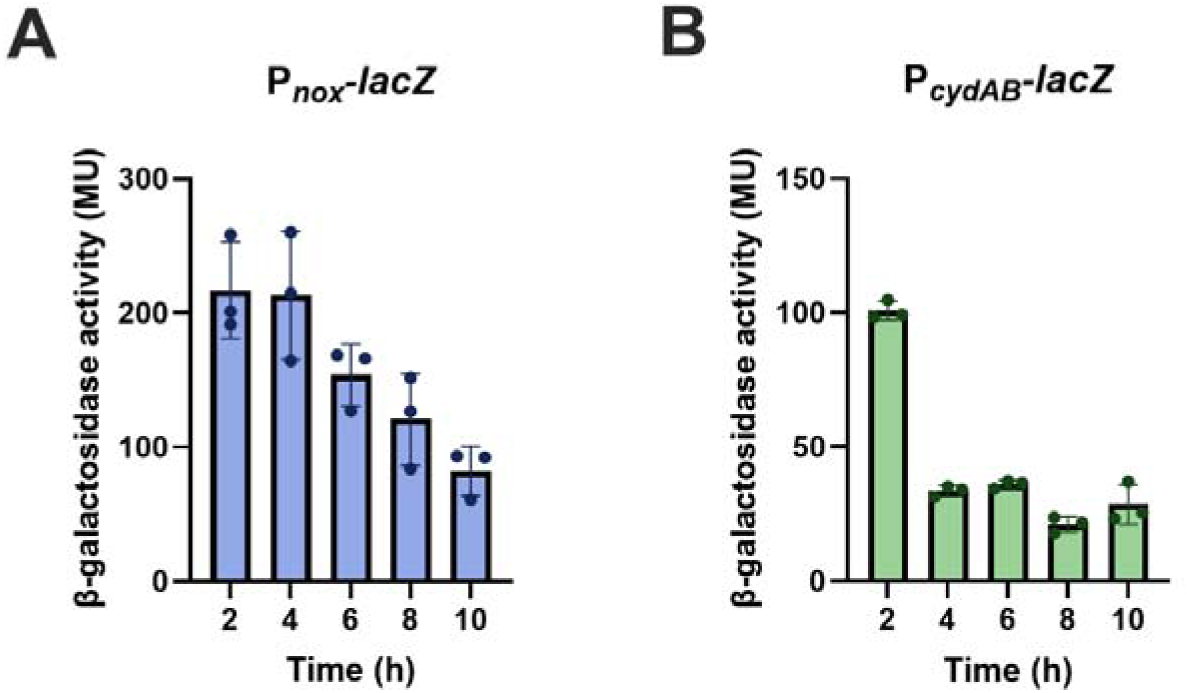
Expression of *nox* and *cydAB* in *E. faecalis* wild-type in hypoxia-permissive conditions over time. *E. faecalis* wild-type expressing *lacZ* reporter constructs P*_nox_*-*lacZ* and P*_cydAB_*-*lacZ* were grown in hypoxia-permissive serum vials for 10 h, with samples taken every 2 h. ß-galactosidase assays were carried out on harvested cells using LacZ activity as a proxy for *nox* (A) and *cydAB* (B) gene expression. The data shown are representative of the mean of biological triplicate ± SD.

**Fig. S3.**
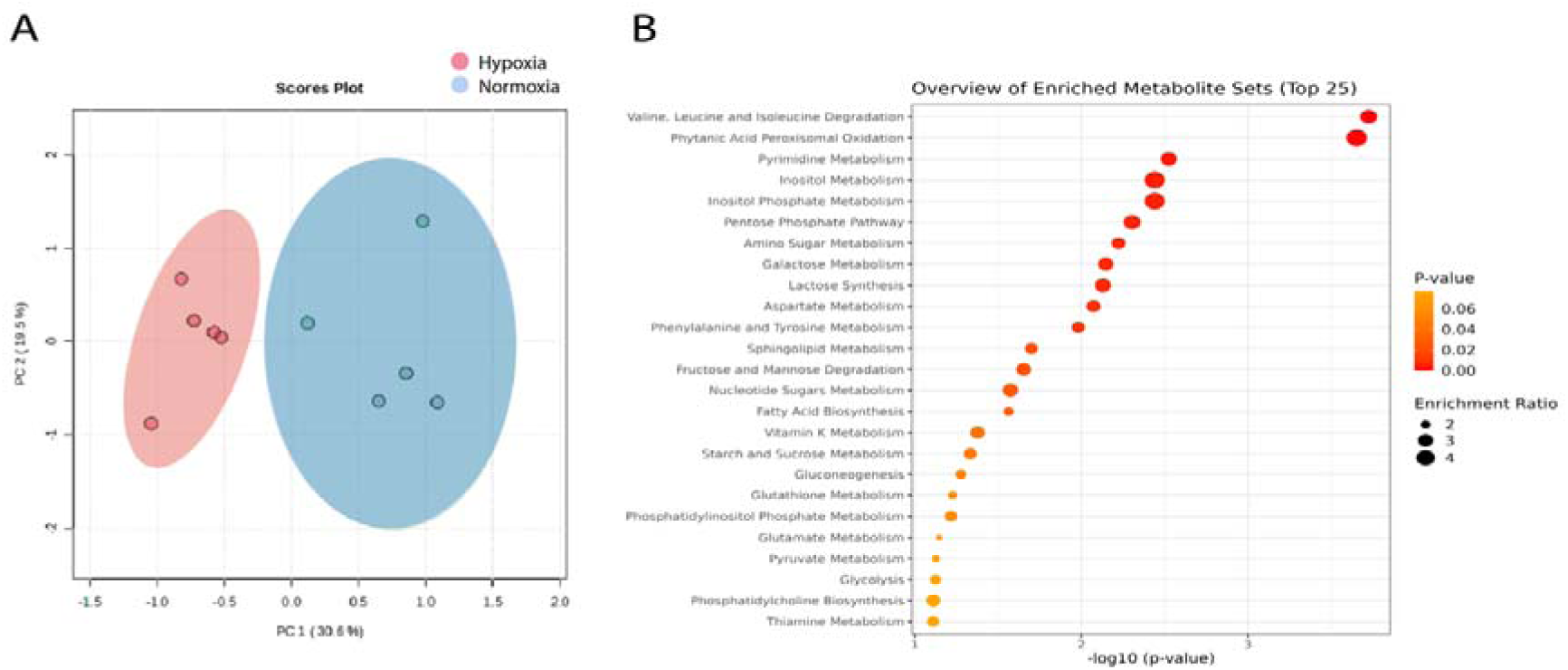
Clustered analysis of the variation between the two distinct adaption groups (hypoxia-adapted and normoxia-adapted) in the wild-type. (A). The principal component analysis of the semi-targeted metabolomic data between the two adapted states (hypoxia and normoxia) in *E. faecalis* wild-type. (B) KEGG pathway enrichment analysis of the metabolites between the two adapted states (normoxia relative to hypoxia) in wild-type. The number of metabolites (from the 130 identified in this dataset) that were enriched in the wild-type hypoxia condition compared to the wild-type normoxia condition for each KEGG pathway was carried out and shown using Metabolanalyst v6.0.

**Fig. S4.**
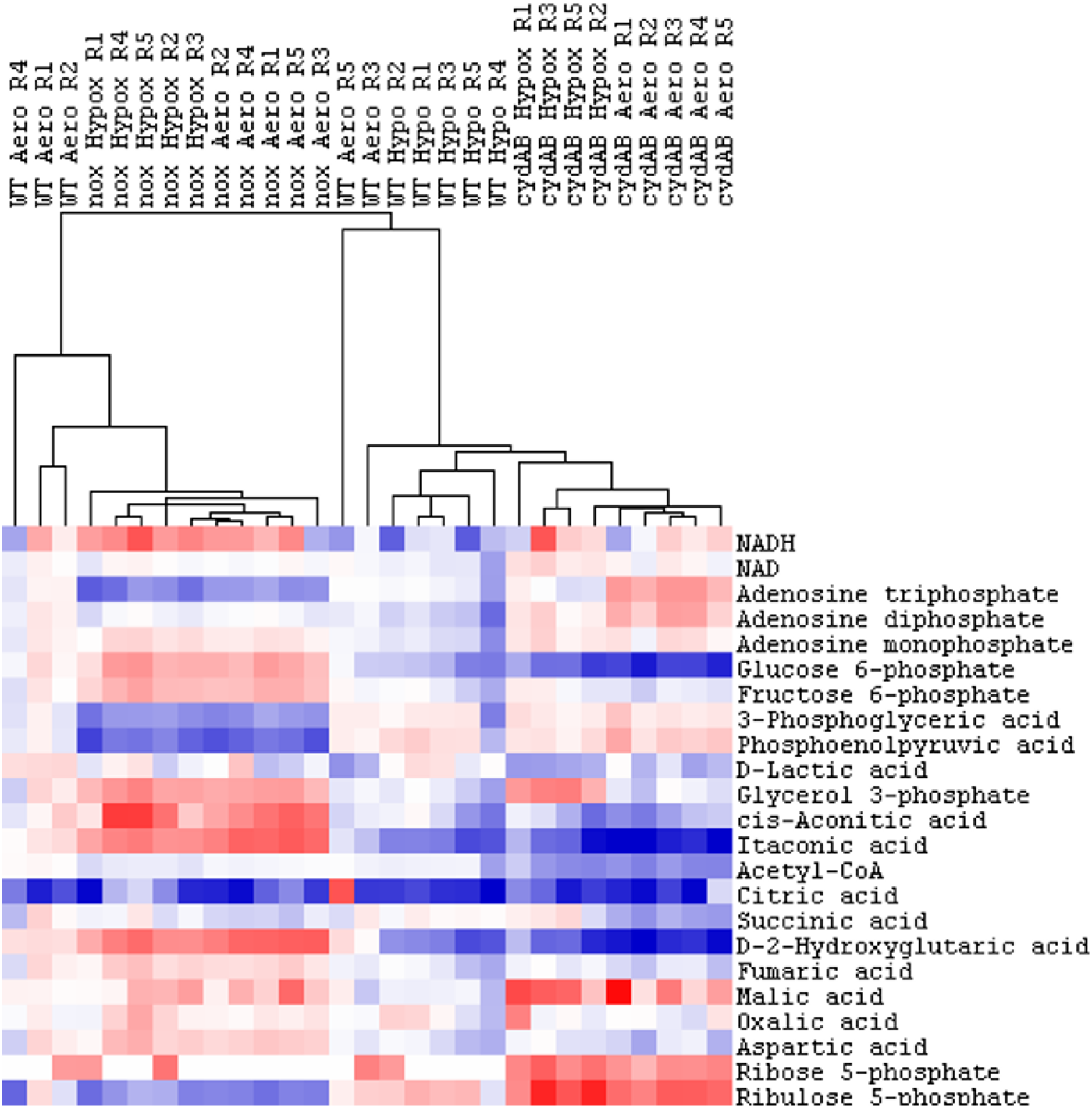
Semi-targeted metabolic clustering of *E. faecalis* wild-type (WT) and respiratory mutants Δ*nox* (nox) and Δ*cydAB* (cydAB) grown under normoxic (Aero) or hypoxic (Hypox) conditions. Each replicate for each strain and each condition is clustered depending on their metabolic profile for the 23 selected metabolites.

**Fig. S5.**
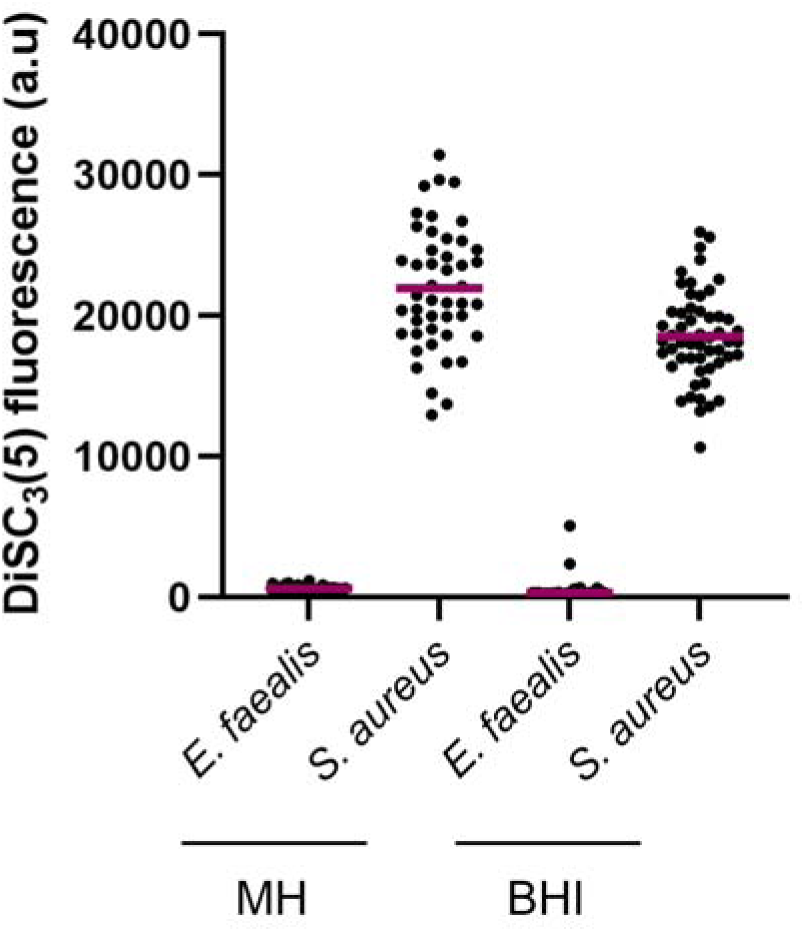
Single-cell measurements of relative membrane potential in *E. faecalis* and *S. aureus*. Cells were grown to early– mid log phase in Brain Heart Infusion (BHI) or cation-adjusted Mueller-Hinton (MH) and stained with the voltage-sensitive dye DiSC_3_(5).

**Fig. S6.**
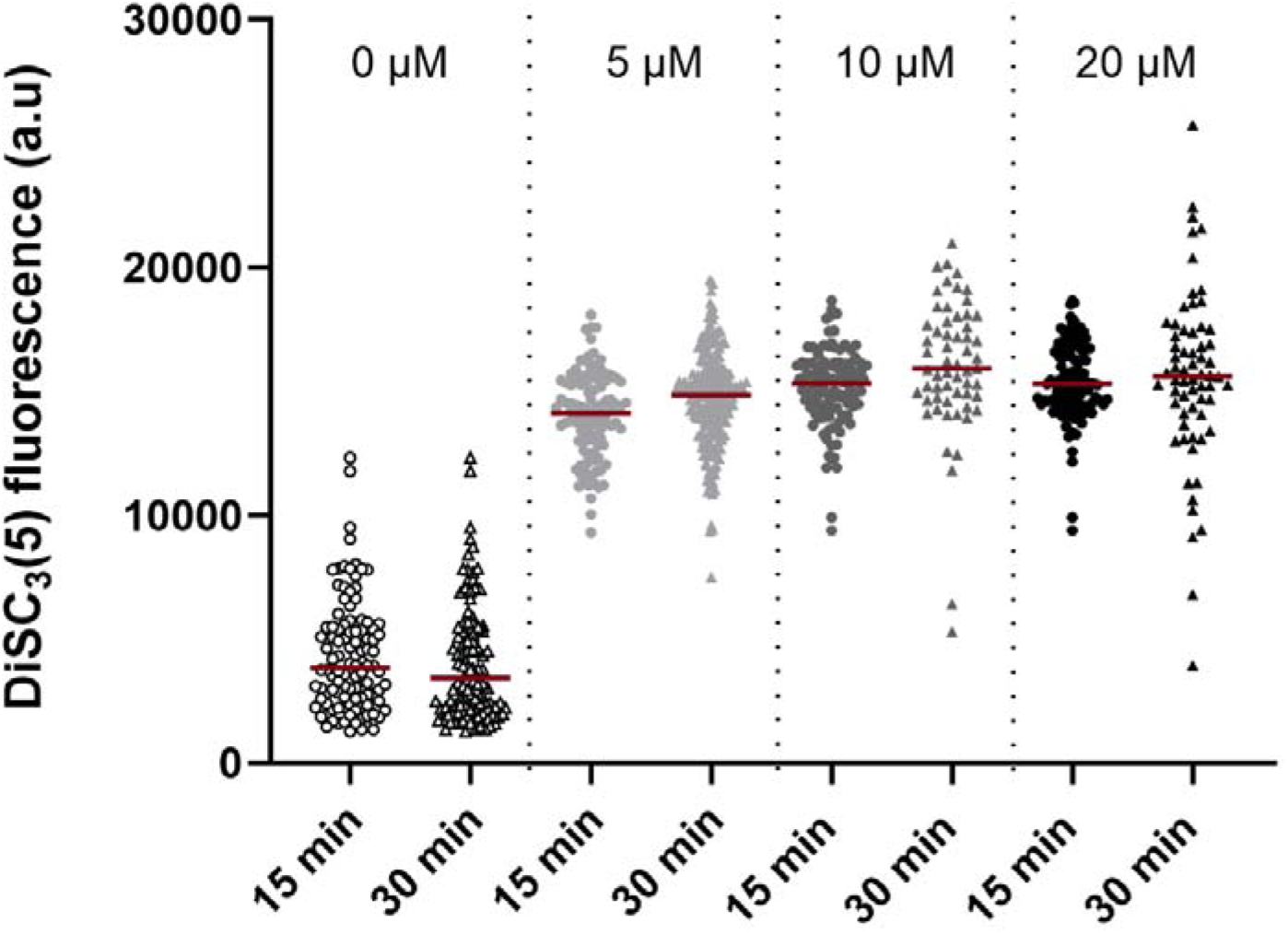
Single-cell measurements of membrane potential in *E. faecalis* wild-type challenged with the protonophore nigericin. *E. faecalis* wild-type was grown to early-mid exponential phase and challenged with nigericin at concentrations of 0, 5, 10 and 20 µM for 15 and 30 min. These data show that the nigericin concentrations and incubation times used were sufficient to achieve steady-state hyperpolarisation. Relative membrane potential were quantified following incubation with the voltage-sensitive dye DiSC_3_(5) and single-cell fluorescence microscopy.

